# Single Photon, Time-Gated, Phasor-based Fluorescence Lifetime Imaging Through Highly Scattering Medium

**DOI:** 10.1101/686998

**Authors:** Rinat Ankri, Arkaprabha Basu, Arin Can Ulku, Claudio Bruschini, Edoardo Charbon, Shimon Weiss, Xavier Michalet

**Affiliations:** Department of Chemistry & Biochemistry, UCLA, Los Angeles, CA; School of Engineering, École polytechnique fédérale de Lausanne, Neuchâtel, Switzerland

**Keywords:** Fluorescence lifetime imaging, Phasor lifetime analysis, Time-gated camera, Single-photon detection, Scattering medium

## Abstract

Fluorescence lifetime imaging (FLI) is a powerful tool for *in vitro* and non-invasive *in vivo* biomolecular and cellular investigations. Fluorescence lifetime is an intrinsic characteristic of any fluorescent dye which, to some extent, does not depend on excitation intensity and signal level. However, when used *in vivo* with visible wavelength emitting fluorophores, FLI is complicated by (i) light scattering as well as absorption by tissues, which significantly reduces fluorescence intensity, (ii) tissue autofluorescence (AF), which decreases the signal to noise ratio and (iii) broadening of the decay signal, which can result in incorrect lifetime estimation. Here, we report the use of a large-frame time-gated single-photon avalanche diode (SPAD) imager, *SwissSPAD2*, with a very short acquisition time (in the milliseconds range) and a wide-field microscopy format. We use the phasor approach to convert each pixel’s data into its local lifetime. The phasor transformation provides a simple and fast visual method for lifetime imaging and is particularly suitable for *in vivo* FLI which suffers from deformation of the fluorescence decay, and makes lifetime extraction by standard fitting challenging. We show, for single dyes, that the phasor cloud distribution (of pixels) increases with decay broadening due to scattering and decreasing fluorescence intensity. Yet, as long as the fluorescence signal is higher than the tissue-like phantom AF, a distinct lifetime can still be clearly identified with an appropriate background correction. Lastly, we demonstrate the detection of few hundred thousand A459 cells expressing the fluorescent protein mCyRFP1 through highly scattering phantom layers, despite significant scattering and the presence of the phantom AF.

## Introduction

Fluorescence microscopy is an invaluable tool in biomedical investigations which holds significant potential for various sensing applications, including probing tissue physiology, detecting early stages of disease *in vitro* and *in vivo* ^*1, 2*^, sensing molecular concentrations of delivered pharmaceutical or intracellular fluorescent proteins^3, 4^. However, its use in the visible wavelength range (400-650 nm) is limited to depths of no more than a few hundred micrometers inside the tissue, due to light scattering and absorption, as well as tissue autofluorescence (AF)^5, 6^. Many of these limitations are mitigated when using near-infrared (NIR)^4, 7^ or short-wave infrared (SWIR) emitting probes^8^, where light scattering and absorption by tissues is minimal, and endogenous fluorophores contribute negligible fluorescence^7, 9^. Nonetheless, *in vivo* fluorescence imaging in the visible range has some advantages: ease of use, low cost, and a large selection of fluorescent dyes with good spectral separation as well as different lifetimes^5, 10^. Conventional *in vivo* imaging methods in the visible range struggle to overcome tissue scattering and tissue absorption^11-13^. A tissue imaging technique based on visible illumination that is simple, fast, compact, portable, versatile and inexpensive is therefore highly desirable.

Fluorescence lifetime imaging (FLI) is a powerful tool for non-invasive *in vitro* and *in vivo* biomolecular and cellular investigations^14-17^. Fluorescence lifetime is an intrinsic characteristic of any fluorescent dye, which, over a broad range of conditions, does not depend on excitation intensity and detected signal level. Moreover, it enables the separation of targeted dyes signal from intrinsic tissue AF, provided their lifetimes differ sufficiently^14^. FLI can therefore provide enhanced sensitivity and contrast while allowing for lifetime-multiplexed detection using dyes with distinct lifetimes.

FLI can be measured in frequency domain (frequency modulated techniques) and in time domain (time-resolved techniques)^18, 19^. Many time-domain FLI techniques use time-correlated single-photon counting (TCSPC), a technique generally used in scanning confocal set-ups^20^ (with exceptions, see e.g. ref.^21^), while the remainder use time-gating methods, which are generally used in wide-field geometries (with no scanning), as in this work. TCSPC methods record the arrival time of each photon after the excitation of a laser pulse, and yield high-resolution histograms of arrival times, while time-gated FLI records the fluorescence decay in a generally smaller number of integration windows (“gates”) of finite width, providing a coarser representation of the fluorescence decay’s temporal profile ^22, 23^.

Wide-field (parallel) data acquisition is highly desirable when imaging live cells or live animals, as it minimizes the temporal delay between data recorded in different regions of the field of view (FOV). It also has the advantage of dispensing with costly and complex scanning devices, even though it creates other challenges. In particular, the laser source needs to be sufficiently intense in order to deliver enough and uniform illumination throughout the whole FOV. In addition, for time-gated acquisition, where the total imaging duration is equal to the product of the number of gates by the integration time per gate, a trade-off between decay resolution (number of gates), signal-to-noise ratio (related to the integration time per gate) and total integration time, needs to be found in order to avoid excessive delay between the start and end of the acquisition.^24^

In this paper, we use a recently introduced wide-field time-gated SPAD camera, *SwissSPAD2*^23^, characterized by low dark-count rate and fully configurable, high-resolution time-gating capabilities, allowing fluorescence lifetime imaging with picosecond time-resolution and acquisition time as low as 2.6 ms per 8 bit gate image. We explore its ability to acquire FLI data through tissue-like phantom, and to enable distinguishing between extrinsic fluorescence and AF, through different phantom thicknesses mimicking skin tissue.

The resulting data were analyzed using the phasor approach^25^, providing a user-friendly graphical representation of fluorescence decays while still allowing quantitative analysis to be performed ^25, 26^. We show that phasor analysis of time-gated fluorescence of visible range dyes, loaded in glass capillaries and imaged through tissue-like phantom layers, can provide the necessary contrast for sub-cutaneous studies. Using the time-gated SwissSPAD2 camera we were able to adjust the acquisition times to the dye concentration, avoiding any saturation or bleaching effects of the excited dye. We use different background analyses and standard deviation analysis to learn how and whether the lifetime changes with the intensity of the fluorescence, compare to the AF of the phantom, as well as with its scattering coefficient. We show that a computational background subtraction results with a correct lifetime for the sample when the fluorescence of the dye shows high intensities, but when the dye shows relatively low concentration, then a phantom BG is critical to extract the correct lifetime. We study the effect of increasing scattering coefficients on the dye’s lifetime SDV and values. Eventually, we show the direct detection of the fluorescent protein (FP) mCyRFP1 expressed in A549 cells through a 1.5 mm phantom layer, enables to distinguish it from control cells.

## Methods

### Wide-field FLI with SwissSPAD2

A schematic description of the experimental setup is presented in Fig. 1(a) capillary (ID: 0.9 mm, OD: 1.2 mm) containing a dye solution (or a solution of suspended cells, see Fig. 1(c), and details below) was held in a custom 3D printed sample holder (see SI for details and CAD drawings). Tissue-like phantom layers (with varying thicknesses of 0.5, 1, 1.5, 2, 3 and 5 mm) were inserted between the capillary and the microscope’s objective lens (x20/0.4, LCPlanFl, Olympus). A 532 nm, 20 MHz repetition rate, <100 ps pulsed laser (LDH-P-FA-530XL, PicoQuant) was used as the excitation source, the incident power (< 12 mW) on the sample being adjusted with the help of neutral density filters. The diffused emitted fluorescence was collected in an epi-illumination mode and imaged on the *SwissSPAD2* (SS2) SPAD array mounted on the side camera port of the microscope. SS2 is comprised of 472×276 single-photon avalanche diode (SPAD)^23^. The in-pixel time-gated architecture affords time-resolved photon-counting at a maximum rate of 97 kfps (1-bit frames). The photon detection efficiency (40 % at 600 nm), fill-factor (10.5 %) and dark count rate (median: 7.5 Hz/SPAD) of the detector is sufficient to collect enough photons for lifetime extraction in low light conditions, such as the case with fluorescence from highly scattering tissue-like phantoms. A finite number (n = 117) of overlapping gate images (gate separation = 428.6 ps) was used, each experiment using a different gate image acquisition time (4 to 400 ms), to account for different sample brightnesses and concentrations. Data files provided in the *Supporting Information* report the number of gates sequence per 1-bit frame acquisition (*exp_no_seq*), from which the total integration time (in ns) can be computed using the following formula:

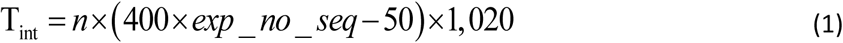

**Figure 1:**
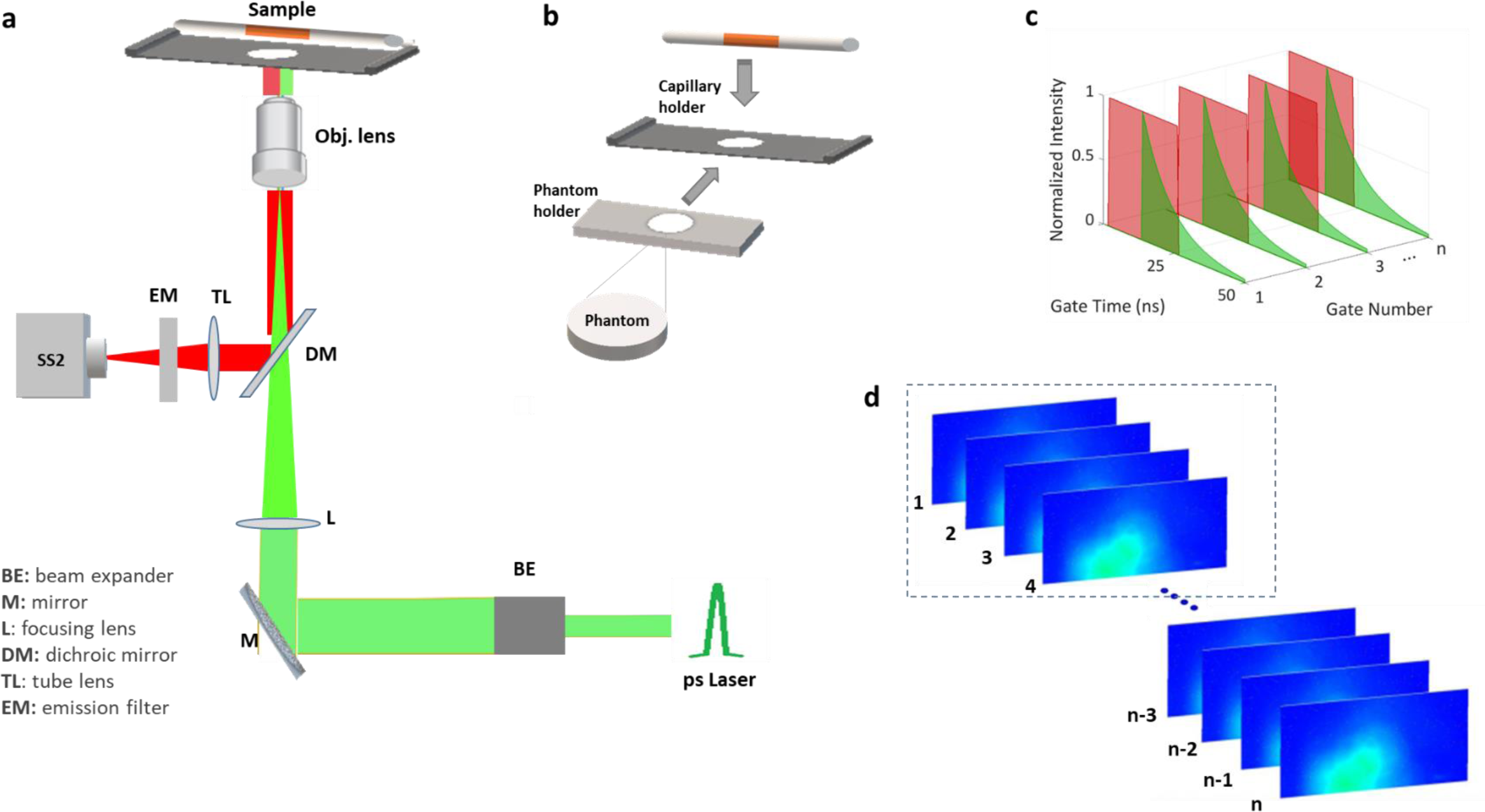
A schematic description of the wide field time-gated FLI setup. (a) FLI set up. The sample was illuminated with a 532 nm picosecond laser and emitted fluorescence was captured using the time-gated Swiss SPAD 2 (SS2) SPAD array (512×256 pixels). Excitation power varied for the different samples, but for all samples behind phantoms we used an incident excitation power of 12 mW. (b) Illustration of the sample preparation procedure. The capillary was deposited on top of a sample holder which held a phantom layer beneath the capillary. The different phantoms layers are characterized by common absorption (µa) and scattering (µs) coefficients but varying thicknesses (between 0.5 to 5 mm). The thickness is defined by that of the 3D-printed phantoms holders. (c) Conceptual illustration of time gating in SwissSPAD2. The delay between subsequent gate positions is a small fraction of the gate width, resulting in overlapping gates. The decay is comprised of 125 such gates. (d) For some samples, multiple series (dashed box) of 125 gate images (n= number of images for all series) were recorded successively, in order to avoid significant bleaching during recording. Variable number of such series were summed post-acquisition to obtain the desired total signal intensity.

Thus, for instance, the shortest value used in these measurements, *exp_no_seq* = 5, corresponds to a total integration time T_int_ = 0.25 s.

### Dyes solutions preparation

Cy3B (GE Healthcare, Waukesha, WI) and ATTO 550 (ATTO-TECH) were prepared with double distilled water (ddH_2_O), with estimated final concentrations of 10 µM (Cy3B) and 1.33 µM (ATTO 550), respectively, and loaded into capillaries (OD: 1.2 mm, ID: 0.9 mm, wall thickness: 150 µm, World Precise Instruments). The total sample volume within each capillary varied between experiments, however, the same FOV dimension was used for all samples and experiments.

### Tissue-like phantom preparation

Solid phantoms were prepared in order to simulate skin tissues with specific optical properties ^27^. The phantoms were prepared using India ink, as the absorbing component and Intralipid (IL) 20% (Lipofundin MCT/LCT 20%, B. Braun Melsungen AG, Germany), as the scattering component^28^. All phantoms contained the same ink (0.003%) and IL (0.75%) final concentrations (v/v), apart for the phantoms used in Fig. 5, which contained increasing concentrations of IL (0.75, 1.5, 2, 4 %). 1% agarose powder (SeaKem LE Agarose, Lonza, USA) was added to the solution to form a gel. Briefly, the solutions were heated and mixed at a temperature of approximately 90 ^°^C while the agarose powder was slowly added. The phantom solutions were stirred continuously to obtain high uniformity. The mixture was then poured into a plastic syringe and stored within water until used. For each experiment, a slightly oversized slice of phantom was cut and placed on the phantom holder comprised of a glass coverslip bottom taped to the 3D printed spacer of appropriate thickness (0.5, 1, 1.5, 2, 3 or 5 mm), cut with a knife to achieve the desired thickness, and covered with another coverslip. This phantom slice was then slid under the sample (capillary) in the 3D printed sample holder assembly.

### Phantom and sample holder

Five phantom holders were designed and 3D printed using the OpenSacd free software, providing five different thicknesses (0.5, 1, 1.5, 2, 3 and 5 mm in thickness). Each phantom holder presented a circled hole in its center, to enable the dye or phantom excitation, as well as their fluorescence collection. A capillary holder (illustrated in Fig 1c) was used to hold the dyes loaded within the capillaries. When a phantom layer was involved in measurement, it was slid under the capillary, after which the latter was first excited with no phantom behind.

### Cells

#### Cell Culture

A549 cells were cultured in Dulbecco’s Modified Eagle’s Medium (DMEM) containing 4.5 g/L D-glucose, 4 mM L-glutamine, 110 mg/L sodium pyruvate supplemented with 10% Fetal Bovine Serum (FBS) and 1% Penicillin/Streptomycin under standard conditions (37 ^o^C and 5% CO_2_).

#### Transfection

Cultured A549 cells were transfected with 2.5 µg of mCyRFP1-C1 plasmid (Addgene, MA) using Lipofectamine 3000 as transfection reagent. 48 hours post transfection, cells were trypsinized, resuspended and centrifuged to collect the cell pellets. The suspensions were then inserted into capillaries, for FLI measurements.

### Phasor analysis

Phasor analysis of the wide-field time-gated FLI data was performed as described previously^29^ with minor modifications required by the presence of detector background and phantom autofluorescence, as described below^30^. All analyses were performed using *Alligator*^*29*^, a freely-available software developed in LabVIEW (see SI for details), dedicated to phasor analysis of time-gated data. Briefly, the uncorrected, uncalibrated phasor (*g*_*i,j*_, *s*_*i,j*_) of each pixel of coordinate (*i, j*) in the image was calculated according to:

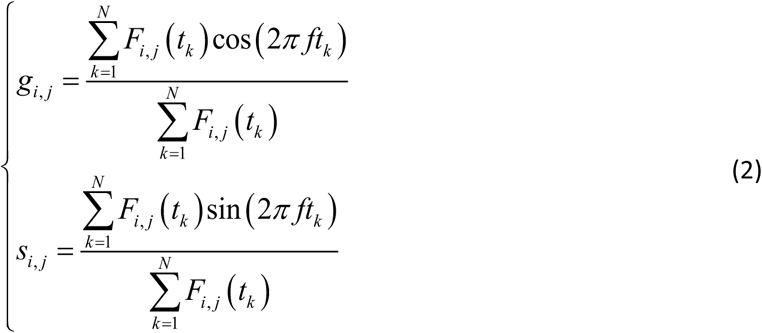

where *f* is the phasor harmonic (chosen in this study equal to the laser repetition rate = 1/*T*), k = 1 … N is the gate number and *F*_*i,j*_(*t*_*k*_) is the k^th^ gate image value at pixel (i,j). When computing region of interest (ROI) phasor values, the *F*_*i,j*_ (*t*_*k*_) in Eq. (2) were replaced by the sum of all *F*_*i,j*_ (*t*_*k*_) in the ROI.

In practice, background-corrected phasors were used as described next.

### Uncorrelated background correction

Although the dark-count rate (DCR) of SS2 is very low (0.18 cps/µm^2^), long integration can result in a significant amount of uncorrelated (i.e. constant) signal added to the contribution of fluorescence. Since the gate duration is constant, this contribution is equal for all gates and can be subtracted to recover the contribution of fluorescence only. Assuming a square-shaped gate of width *W* (a good approximation in these experiments}) and a single exponential decay with amplitude *A* and lifetime *τ* (an approximation whose validity depends on which sample is considered), the uncorrelated background contribution *B* can be estimated from the minimum and maximum recorded gate intensities (*F*_*min*_ and *F*_*max*_, recorded for gate *k*_*min*_ and *k*_*max*_, which are easily identified graphically), as well as the median gate intensity (*F*_*med*_, recorded for gate *k*_*med*_ = ½ (*k*_*min*_ and *k*_*max*_)):

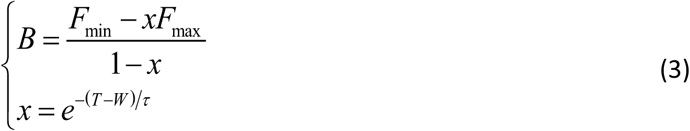

where *x* is the solution of a quadratic equation with solution:

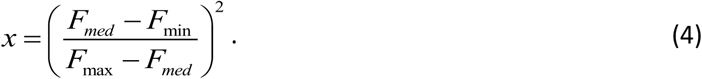

The amplitude of the background-subtracted decay is equal to:

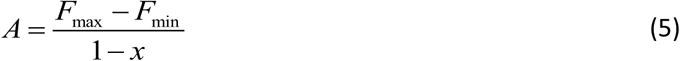

and the integrated background-subtracted signal (which can also be calculated by subtracting *B* from the decay and summing up the corrected gate signals) is given by:

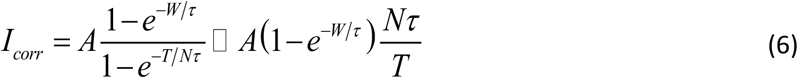

Note that an estimate of the decay’s lifetime can readily be obtained using *x*:

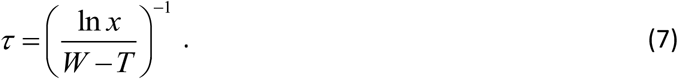

Using the background-corrected gate intensities *F*_*i,j*_(*t*_*k*_) − *B* instead of *F*_*i,j*_(*t*_*k*_) in Eq. (2) results in the uncorrelated background-corrected phasor.

### Phantom autofluorescence correction

In the presence of a known phantom autofluorescence background (measured by recording the phantom in the absence of sample, in the exact same conditions of excitation intensity and integration time,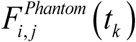), an autofluorescence (and uncorrelated background) -corrected phasor can be computed, by replacing *F*_*i,j*_(*t*_*k*_) by 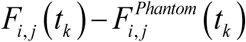in Eq. (2).

### Phasor calibration

Background-corrected phasors computed as discussed still need to be calibrated using a reference sample with known lifetime *τ*_*c*_, in order to account for the finite instrument response function (IRF)^29, 31^, as discussed next.

The theoretical phasor of the single-exponential decay calibration sample (located on the UC) is given by^25,32^:

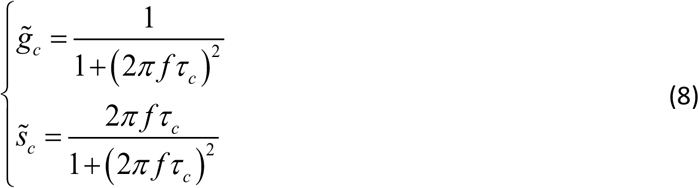

Equivalently, treating the phasor (*g, s*) as a complex number *z*:

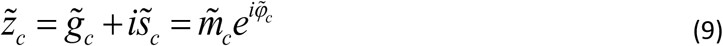

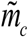 and 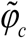 are the calibration phasor’s modulus and phase, respectively, given by:

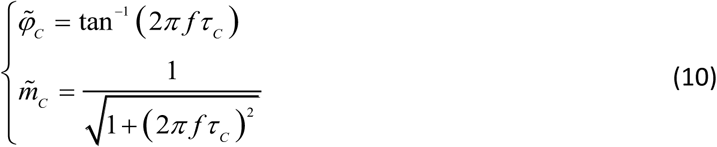

The reason for introducing these quantities is that the effect of a finite IRF on the phasor of any decay is to convolve the “pure” decay with the IRF, which, in the phasor representation, amounts to a simple multiplication by the IRF’s complex phasor.

Calling *z*_*c*_ the measured (uncalibrated) phasor of the reference sample, the calibrated phasor 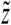 of any measured sampleis then given by:

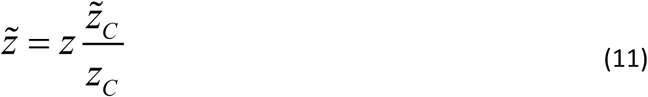

or, using the modulus and phase notation:

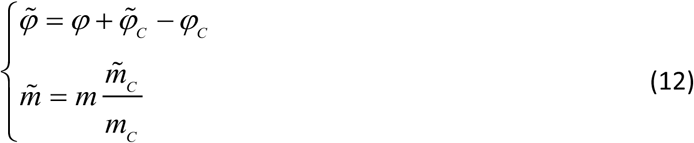

### Phase lifetime dispersion

For any given sample (even one which is not characterized by a single exponential decay), it is possible to compute a “phase lifetime” *τ*_*φ*_ from the calibrated phasor by inverting the first expression in Eq. (10):

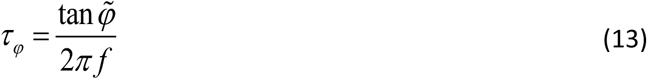

(A “modulus lifetime” can also be computed using the inverse of the second expression in Eq. (10), but its usefulness is limited by the fact that it requires 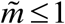, which is sometimes false due to noise).^25^

In order to study the respective contribution of signal intensity and scattering on the phase lifetime dispersion, a plot of phase lifetime versus intensity for a given sample is needed. This can be obtained in one of two ways:

- Several acquisitions with homogeneous intensity within the FOV, but recorded with different integration times.
- A single acquisition with inhomogeneous intensity within the FOV.

For samples observed behind a phantom layer, intensity inhomogeneity within the FOV is limited due to scattering, and therefore the first approach is necessary. For samples observed without phantoms, the non-uniformity of the illumination spot naturally creates fluorescence inhomogeneity throughout the FOV, and the second approach is the most efficient.

In either case, given a sample time-gated dataset, phase lifetimes{*τ*_*φ,i*_}, *i* = 1 … *R* of non-overlapping 4×4 pixel regions of interest (ROIs) are calculated and a scatter plot of *τ*_*φ,i*_ versus *I*_*i*_, the total intensity of ROI *i* is computed.

In the first approach described above, the mean intensity of all ROIs, *Ī*, is computed, while the histogram *h*(*τ*_*φ*_) of {*τ*_*φ, i*_} is fitted with a suitable function. We found out that, due to the asymmetry of this distribution, a bi-Gaussian function is adequate, provided a robust fit rejecting outlier is used:

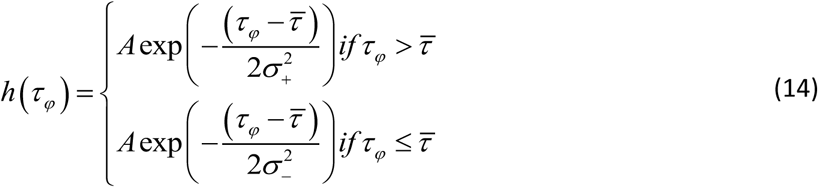

The standard deviation *σ* of this function is given by:

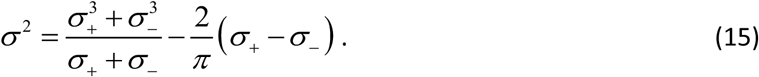

This standard deviation is used as the measure of the phase lifetime dispersion for that average intensity. Repeating this operation with a sample of varying mean intensity *Ī*, we obtain a function *σ*(*Ī*) describing the dependence of the phase lifetime dispersion on average intensity.

In the second approach (non-uniform intensity throughout the FOV), the same type of scatter plot of *τ*_*φ,i*_ versus *I*_*i*_ is computed, and then divided in “data slices” characterized by *k*Δ*I* ≤ *I*_*i*_ < (*k* +1) Δ*I*, where Δ*I* is the slice’s width, chosen such that each slice contains a sufficient number of data points to compute a meaningful standard deviation of the corresponding phase lifetimes (e.g. > 100 data points). We thus obtain, with a single sample dataset, a relation *σ* (*Ī*_*k*_), where 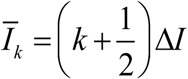 is the mean intensity of slice *k* as illustrated in Fig. 2d.

**Figure 2:**
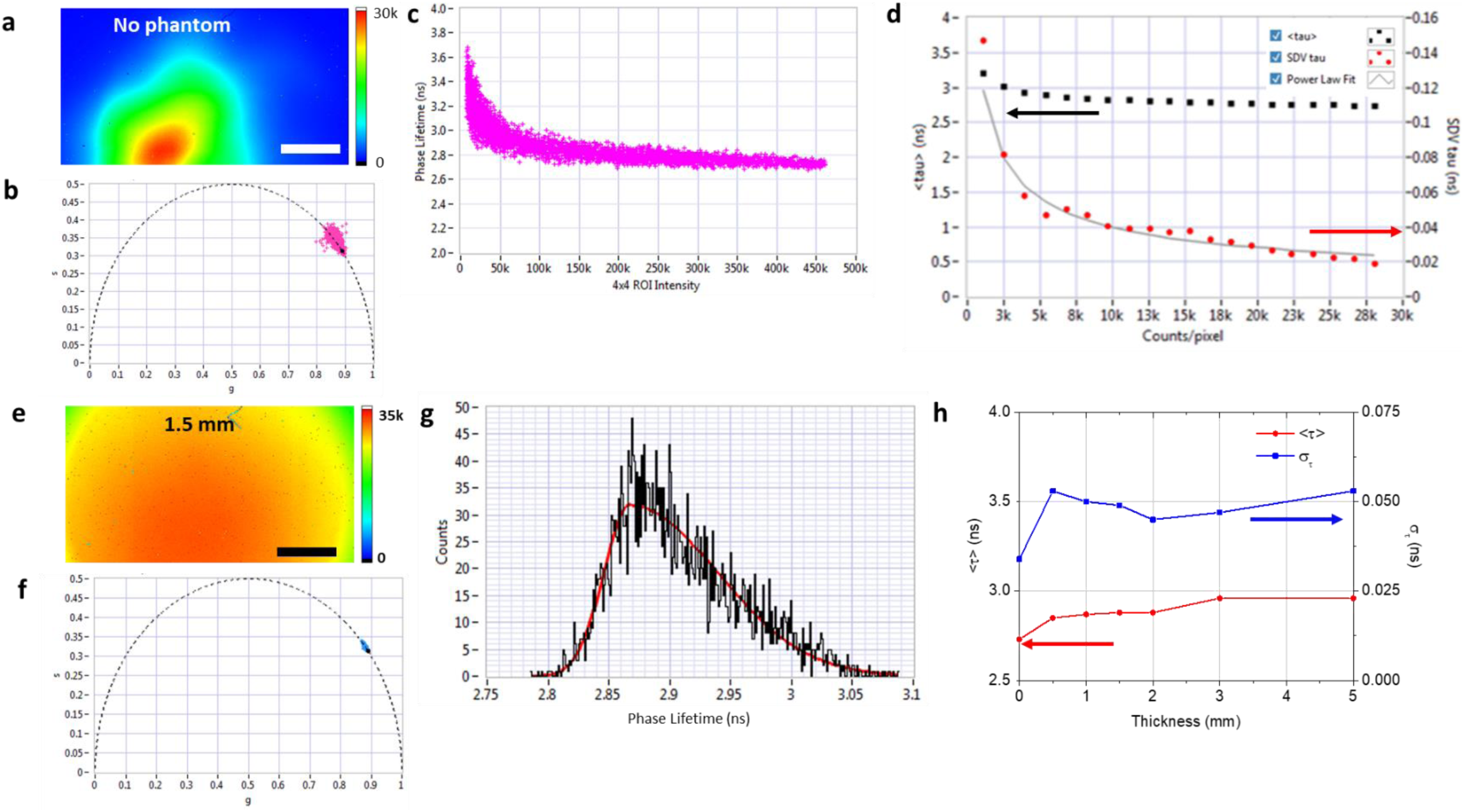
Phase lifetime analysis of Cy3B behind different phantom thicknesses. (a) & (e): Color-coded intensity images of the Cy3B sample observed behind no phantom or a 1.5 mm phantom, respectively. € shows a more homogeneous fluorescence intensity, due to scattering in the phantom. Scale bar is 160 µm. (b) & (f): Phasor scatter plots for the datasets shown in (a) and (e), respectively, calculated using 4×4 pixel ROIs covering the lower half of each image (see Figure S1a in *Supporting Information* for details). The phasors are calibrated using the average phasor of the bottom half of the dataset shown in (a) (*Supporting information* Figure S1b) as calibration, using the known Cy3B lifetime (τ = 2.8 ns) and a phasor harmonic frequency f = 20 MHz. (c) Scatterplot of phase lifetime versus total intensity calculated for the phasors shown in (b). (d) Analysis of the dependence of mean phase lifetime <τ> and standard deviation στ on total intensity (expressed in counts/pixel). The latter is well described by a power law with exponent α = −1/2, consistent with shot noise-limited measurement. (g) Phase lifetime histogram corresponding to the phasor plot shown in (f) and fit with a bi-Gaussian (red curve), from which a peak phase lifetime and standard deviation are extracted. (h) Peak phase lifetime and SDV of the 10 µM Cy3B sample, computed as described in (g), as function of phantom thickness. For all samples, the average fluorescence intensity per pixel was 30,000 counts. The phase lifetime SDV of the Cy3B sample observed without phantom was obtained by extrapolating the curve obtained in (d).

No matter which approach is used to compute this relation between phase lifetime dispersion and mean intensity, an inverse square root function 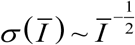 indicates that the phase lifetime dispersion is dominated by shot noise, while any departure from this behavior suggests that additional effects are at play.

## Results and Discussion

### Effect of increasing phantom thicknesses on the phase lifetime

We first performed time-gated phasor FLI analysis of a bright Cy3B sample loaded in a thin glass capillary (concentration: 10 µM) placed behind various phantom thicknesses, in order to study the influence of scattering on the recorded signal intensity and measured fluorescence phase lifetime. Figure 2a shows the fluorescence intensity image of the Cy3B sample in the absence of phantom, as recorded with SS2 using an excitation power of 0.5 mW (measured before the objective lens), and a 0.25 s integration time. A scatter plot of phasor transformations of 4×4 pixel ROIs (see *Supporting information*, Fig.S1a) is shown in Fig. 2b, after uncorrelated background (ucB) correction using Eq. (3)-(7) and calibration with the average phasor of the sample using the literature lifetime value τ = 2.8 ns (see *Supporting information*, Fig. S1b). The corresponding phase lifetimes (Eq. (13)) are plotted versus the 4×4 ROI intensities in Fig. 2c, indicating that the phase lifetime variance increases for ROIs of lower intensity, while the phase lifetime itself shows a slight positive bias at low intensities. Fig. 2d presents the phase lifetime standard deviation (SDV) as a function of mean intensity per pixel obtained from the data shown in Fig. 2c, fitted with a power law:

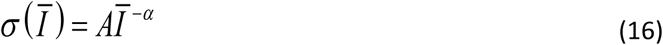

where *α* = 0.50 ± 0.03 as expected for a shot noise limited observable ^31^.

The same analysis (using the same calibration phasor) was then performed on the sample illuminated and observed through phantom layers of various thicknesses, to simulate measurement through various amounts of tissue. In order to compensate for the attenuation of excitation and emission intensities due to absorption and scattering, increasing integration times were used, in order to obtain comparable total collected signal for all measurements^33^. As an example, Fig. 2e shows the image of the Cy3B sample observed behind a 1.5 mm thick phantom layer. Because of the increased integration time, the level of ucB increased proportionally (*Supporting information*, Fig. S2a) and had to be subtracted to recover the residual fluorescence signal. To do so, we used Eqs. (3)-(7), which assume that the recorded signal is described by a single exponential decay integrated over a square gate. As we shall see later, this assumption is appropriate in this particular experiment, but needs not be in general. Once corrected for ucB and normalized by the integration time, the data yields a recorded intensity which exhibits an exponential dependence on phantom thickness, with a characteristic length scale *d*_*0*_ = 0.77 ± 0.27 mm (see *Supporting information*, Fig. S3b)^23^ (which is close to the scattering coefficient of the sample, 0.75 mm^-1^).

The increase in the phantom thickness increased the scattering events of the photons, resulting with a more homogeneous collected intensity over the whole FOV. Correlatively to this reduced intensity variance, the phasor variance was also reduced (Figure 2f) and consequently, so was the phase lifetime variance (Fig. 2g)^31^.

To disentangle the influence of intensity variance on phase lifetime variance, we compared measurements with different phantom thicknesses (including no phantom) and different integration times but resulting in the same averaged pixel intensity. If both phantom thickness and signal level both contributed to the variance, fixing the intensity would allow isolating any contribution to phantom thickness only. The result of this analysis is presented in Fig. 2h, which shows phase lifetime (red) and phase lifetime’s SDV (blue) for the same Cy3B sample as a function of phantom layer thickness. The measured phase lifetime is essentially constant (2.90 ±0.05 ns), the small positive bias observed with increasing thickness being most likely due to phantom AF (see below), a phenomenon also observed for low signal values in Fig. 2c. The phase lifetime’s SDV is fairly constant for measurements behind phantoms (*σ*_*τ*_ = 0.050 ± 0.03 ns), and slightly larger than that observed in the absence of phantom (*σ*_*τ*_ = 0.034 ns). This difference of the phantom-less sample is most likely due to the spatially inhomogeneity of the signal, with very bright pixels that could result in lower shot noise. Both results (constant phase lifetime and phase lifetime variance) are the phasor translation of the time domain observation, resulting in a measured decays insensitive to the amount of phantom material placed between the excitation source and sample, and between sample and detector (*Supporting information*, Fig. S2).

Importantly, as described in the Methods section, all the above mentioned lifetime calculations were performed using a ‘global phasor’ calibration, i.e., the average phase lifetime of the Cy3B (no phantom) sample. In order to check whether a calibration method taking into consideration local timing differences existing between different pixels, we also used a local calibration method, where the phasor of each 4×4 pixel ROI of the Cy3B (no phantom) sample to calibrate the corresponding ROI of the sample of interest. There was no significant change to the phasor SDV, as well as the FLT, resulting in values very close to those obtained using a global phasor calibration method (data not shown).

### Phantom autofluorescence

To investigate the possibility that the source of the observed phase lifetime bias for thick phantom is due to phantom AF, we performed similar measurements in the absence of sample. Fig.3 shows the result of this analysis, using similar integration time (24 s) and identical excitation power for all phantom thicknesses. The phantom’s AF phasors appear fairly well represented by a single exponential decay with lifetime *τ*_*ph*_ = 3.44 ns superimposed to the ucB contribution of the detector (Fig. 3a,b). The standard deviation of the phase lifetime also appears insensitive to the phantom’s thickness (Fig. 3c), consistent with the result obtained for the Cy3B sample observed through these different phantom thicknesses. Interestingly, the measured ucB-corrected count rates of all these phantoms are comparable (∼ 1 kHz, Fig. S3a) despite their very different thicknesses, suggesting that only a superficial layer (< 0.5 mm) of the phantoms contributes most of the observed phantom AF. This observation is consistent with the fact that excitation and emission intensities are attenuated by absorption and scattering at larger depth.

**Figure 3:**
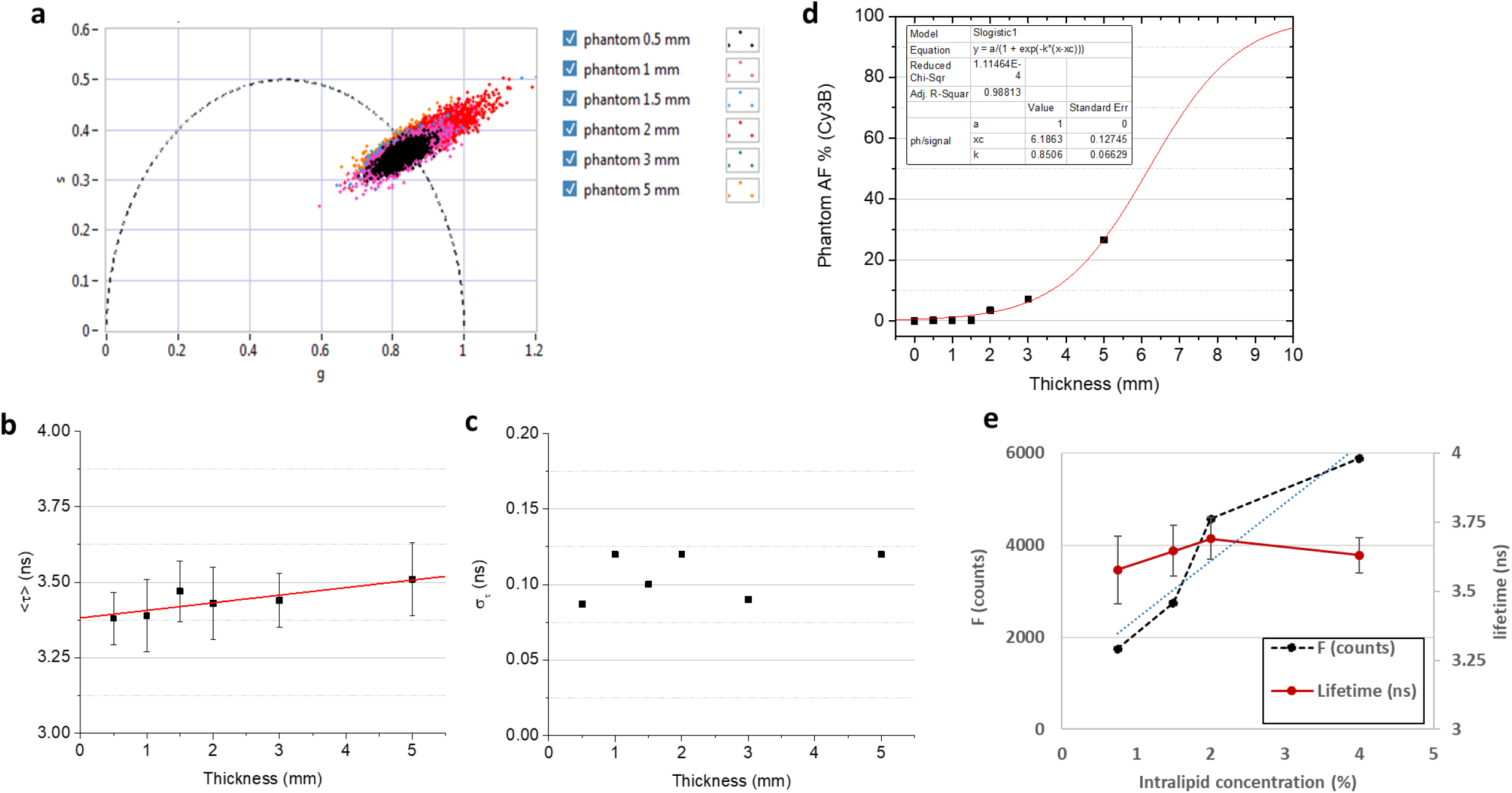
Phantom phasor analysis. (a) Phasor plot of the 6 phantoms. The dispersion along a line connecting the origin and the phasor of a single exponential decay with lifetime τ = 3.44 ns is due to over- or under-correction of the uncorrelated background expected for these low intensity signals. (b) The average phase lifetime measured for all thicknesses is very weakly dependent on thickness (increasing by 25 ps/mm). Error bars are the standard deviations calculated by bi-Gaussian fitting of the phase lifetime histograms, as explained in Fig. 2g. (c) The phase lifetime’s standard deviation is also essentially constant, a translation in the time domain of the similarity of the scatter plots shown in (a). (d) The measured phantom autofluorescence count rate (∼ 1 kHz) can be used to estimate its contribution to the Cy3B signals studied in Fig. 2. Its contribution is negligible below 2 mm, but reaches 27% for the 5 mm phantom. The model fitted to the data is discussed in the text. (e) AF phase lifetime (solid line) and intensity (dashed line) for 0.5 mm phantoms with different intralipid concentrations. The AF increases linearly (dotted line) with the intralipid concentration, which is accompanied by a decrease of the phase lifetime SDV.

With this information in hand, it is possible to revisit the Cy3B data and analyze the influence of phantom autofluorescence on the results shown in Fig. 2. Indeed, because the fluorescence count rate decreased with increasing phantom thickness, integration time was increased to obtain a similar total signal for all samples, thus increasing the fractional contribution of the phantom autofluorescence to the total signal. Fig. 3d represents this fraction, calculated as:

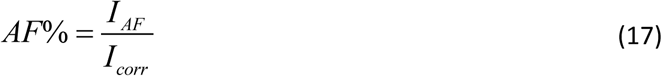

where *I*_*AF*_ is the phantom ucB-corrected autofluorescence count rate (reported in *Supporting information*, Fig. S3a), and *I*_*corr*_ is the ucB-corrected count rate (*Supporting information* Fig. S3b). The contribution of phantom autofluorescence increases steadily with phantom thickness, due to the reduction of the Cy3B sample’s count rate, reaching 27% for the 5 mm phantom. The evolution of AF% is well fitted by a model assuming a constant contribution of phantom AF (rate *I*_*AF*_) and an exponential dependence of the sample fluorescence count rate on phantom thickness, *d*:

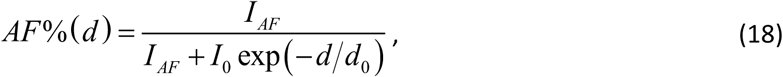

with *d*_*0*_ = 1.18 ± 0.09 mm. Encouragingly, despite phantom AF, the phase lifetime of the Cy3B sample observed behind this same phantom is not markedly changed compared to measurements performed behind thinner phantoms (Fig. 2), most likely because of the large difference between Cy3B’s lifetime (2.8 ns) and that of the phantom’s autofluorescence (∼3.4 ns).

To further point the likely source of phantom AF, we studied phantoms of identical thickness (0.5 mm) but comprised of different intralipid (IL) concentrations. The signal increased linearly with concentration (Fig. 3e), with a small residual offset at 0 % IL pointing at a minimal contribution of the remaining ingredients constituting the phantom (agar, India ink). The AF phase lifetime did not depend on the IL concentration, however, while its SDV decrease as the total amount of AF increased, as expected (Fig. 3f). Since IL is used as a scattering agent, this AF component is an unfortunate side effect, which could possibly be alleviated using a different scattering component.

### Testing the limits of phase lifetime measurements behind phantoms

To test whether phantom autofluorescence could affect the measured phase lifetime of other samples with lifetimes closer to that of phantom autofluorescence or with lower brightness, we looked at a dim sample of ATTO 550 (*τ* = 3.6 ns). Fig. 4a shows the phase lifetimes extracted from a series of measurement of the same ATTO 550 sample measured behind increasing phantom thicknesses, using the same integration time for all acquisitions. We compared two approaches for this calculation: (i) the approach used for Cy3B, using ucB-correction of the signal using Eqs. (3)-(7), which assume that the recorded signal is described by a single exponential decay integrated over a square gate (red data points) and neglects the presence of phantom AF, and (ii) ucB and AF correction by direct subtraction of the phantom-only signals obtained in the series of experiments described in the previous section (blue data point). In both approaches, the same trend is observable: for large phantom thickness (*d* > 3 mm), as the count rate due to ATTO 550 fluorescence decreases (Fig. 4c, blue curve), the extracted phase lifetime becomes indistinguishable from that of the phantom (Fig. 4a, black data points, reported from Fig. 3b). The analytical ucB subtraction approach (red curves), neglecting the presence of the phantom AF, is more sensitive to the phantom thickness as evidenced by the continuous decay of the calculated phase lifetime for thicker phantoms (Fig. 4a, red curve), while the effect of phantom AF only manifests itself for the 5 mm phantom in the case of direct phantom signal subtraction (blue curve). Overall, it appears that as long as the fluorescence signal is at least twice as large as the phantom AF (Fig. 4c), the lifetime of that sample can be distinguished fairly robustly (see also in *Supporting Information* Fig. S5), even when it is close to the phantom AF lifetime (ATTO 550: 3.6 ns, AF: 3.44 ns).

**Figure 4:**
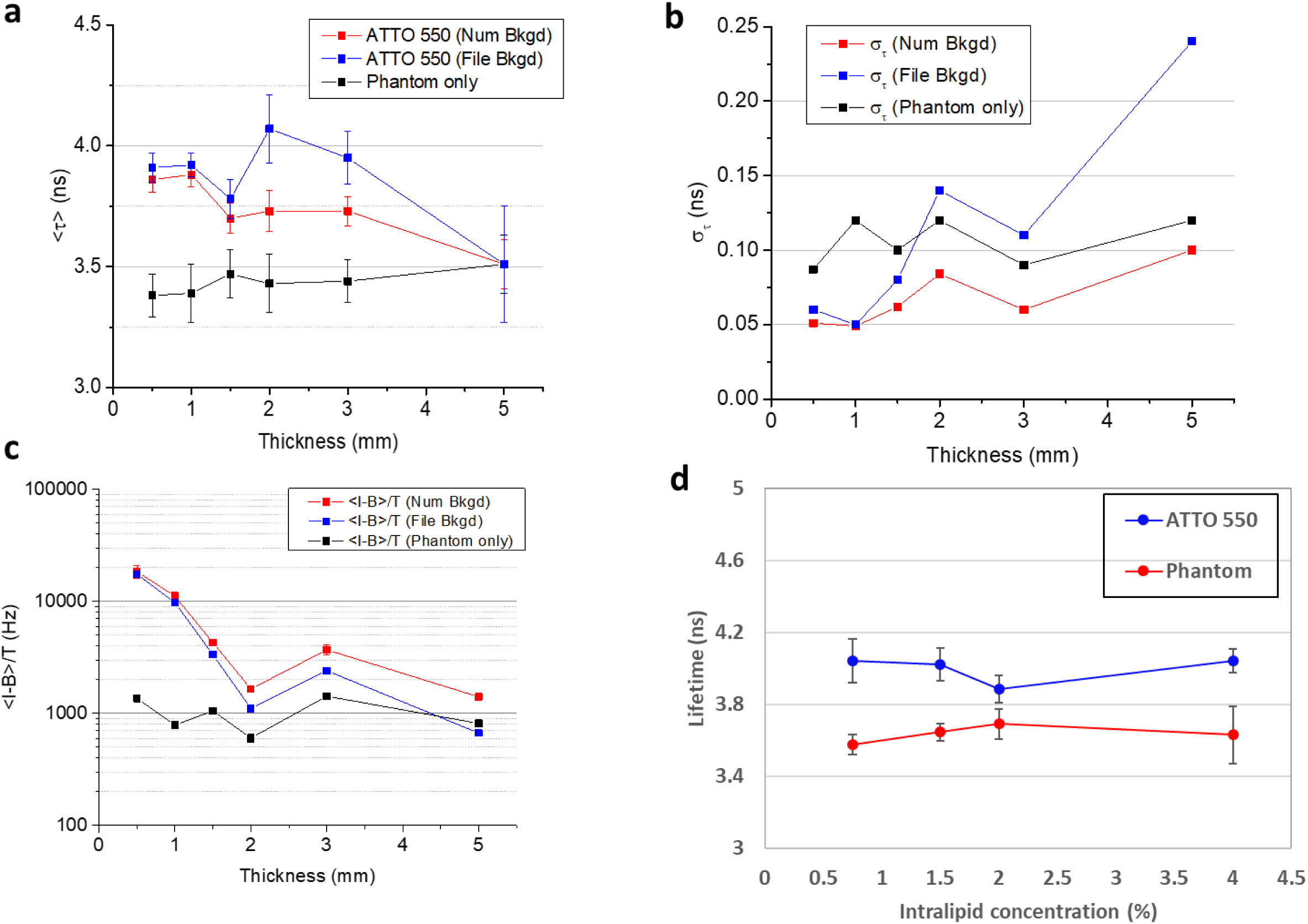
ATTO 550 phase lifetime analysis. (a) Phase lifetime of the ATTO 550 sample observed behind phantoms of different thicknesses, calculated by estimating the uncorrelated background numerically (red curve) or by subtracting the phantom only signal (blue curve). The black curve shows the phase lifetimes of the pure phantom samples. Error bars are the standard deviations calculated by bi-Gaussian fitting of the phase lifetime histograms, as explained in Fig. 2g. (b) Phase lifetime standard deviation (SDV, shown in (a) as error bars) as a function of phantom thickness. The phantom-only phase lifetime SDV (black curve) is essentially constant. The SDV of the phase lifetime calculated by numerical estimation of the uncorrelated background (red curve) doubles from 0.5 mm to 5 mm, while that estimated by phantom-only data subtraction (blue curve) quadruples. (c) Count rate (per 4×4 pixel ROI) as a function of phantom thickness. The phantom AF (black curve) is practically independent from the phantom thickness, while the sample’s fluorescence decreases exponentially with increasing thickness. In the absence of AF correction (red curve), the total signal used for phase lifetime calculation is larger, explaining the smaller standard deviation shown in (b). Error bars indicate standard deviation. (d) ATTO 550 (blue) and AF (red) phase lifetime behind 0.5 mm phantom layers with increasing intralipid concentrations (0.75, 1.5, 2 and 4%). ATTO 550 phase lifetime calculations were performed using phantom layer AF subtraction, using the same intralipid concentration and integration times (10 ms) for each sample. Using AF subtraction, ATTO 550’s measured phase lifetime does not depend on intralipid concentration. However, the SDV depends on signal intensity, as expected for a shot noise limited signal.

**Figure 5:**
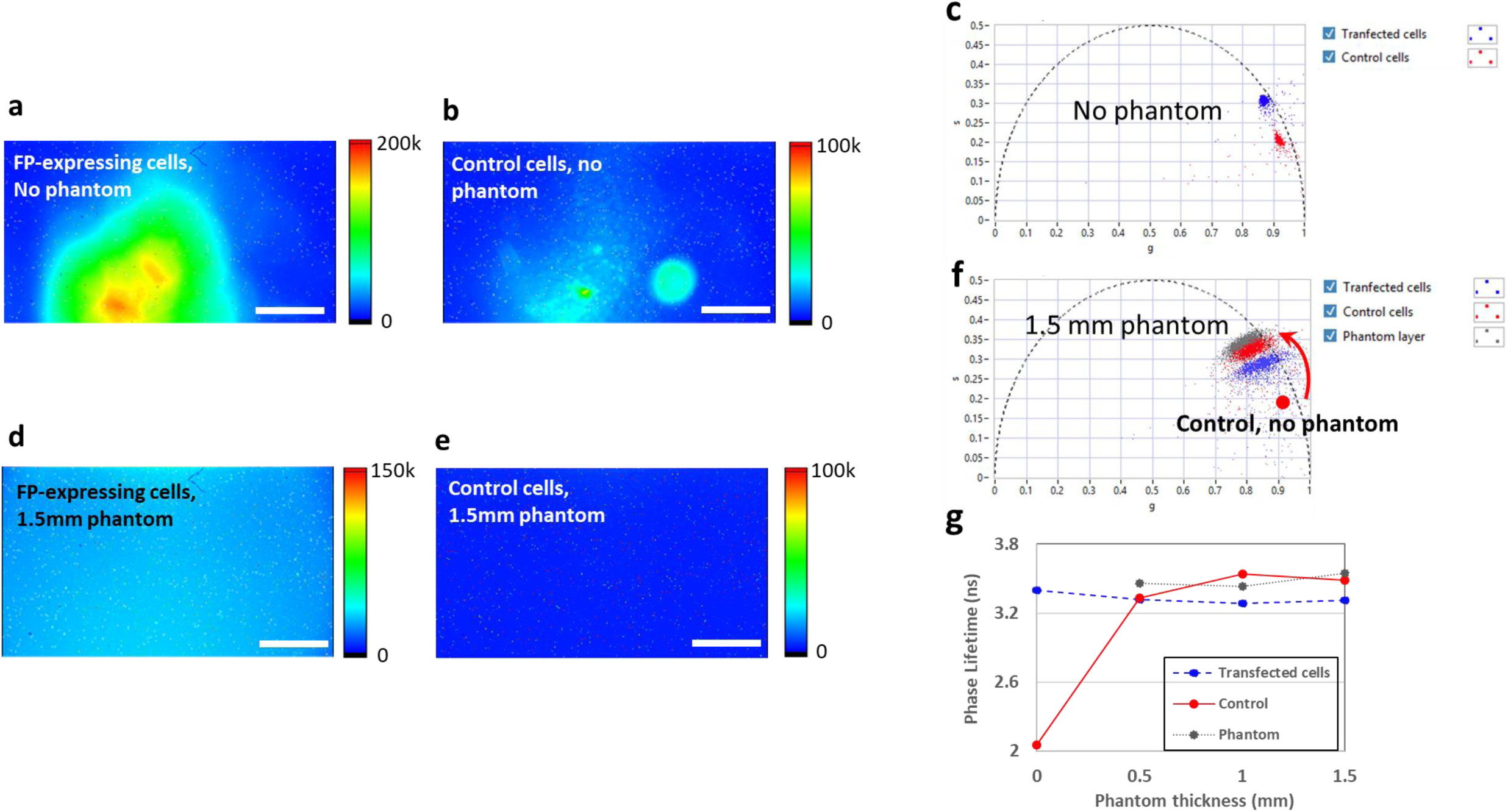
Fluorescence phase lifetime analysis ofA549 cells expressing the fluorescent protein mCyRFP1 behind phantom layers of 0.5, 1 and 1.5 mm in thickness. (a,b): Fluorescence images for A549 cells expressing mCyRP1 and control cells, without phantom. (c) Phasors scatter plots for control and transfected A459 cells without phantom, showing two well distinguishable lifetimes of 2.05 and 3.4 ns, respectively. (d,e) Fluorescence image of A549 expressing mCyRP1 and control cells behind a 1.5 mm phantom. The fluorescence intensity is much lower (a mean intensity of ∼40,000 counts) compared to the mean fluorescence intensity of these cells with no phantom shown above (∼120,000 counts). For all images, scale bar is 150 µm. (f) Phasors scatter plots for A459 cells without phantom and behind a 1.5 mm phantom. A marked jump of the apparent phasor of untransfected cells is noticeable. The resulting phasor is very close to that of the phantom, whose autofluorescence signal is dominating the cells’ autofluorescence. By contrast, the phasor of the transfected cells remains essentially unchanged. (g) Phase lifetime of A549 cells expressing mCyRP1 (dashed line), control cells (A549, which did not express the FP, solid line) and phantom layers AF (0.5, 1 and 1.5 mm thickness, dotted line). The sample containing the control cells (AF phase lifetime: 2.1 ns) has a phase lifetime which is indistinguishable from that of the phantom AF as soon as it is observed behind any amount of phantom, while the cells expressing a FP exhibited a constant phase lifetime of 3.3 ns, no matter what phantom thickness it was observed behind. Analysis was limited to the bottom quarter of the images as detailed in SI.

The measured phase lifetime standard deviation (Fig. 4b), which can be used as a measure of how different the AF and sample lifetimes need to be in order to be distinguishable, depends on the phantom AF: without AF subtraction (red curve), the standard deviation only doubles from 0.5 mm to 5 mm, while when subtracting phantom AF, the measured phase lifetime standard deviation quadruples. This behavior mirrors the different in total count rate measured in the two cases (Fig. 4c): when including phantom AF (red curve), the fluorescence count rate only decreases by a factor 13 from 0.5 mm to 5 mm, while the AF-subtracted count rate (blue curve) decreases by a factor 25.

We performed the same measurements with the 0.5 mm phantoms with varying IL concentration studied in Fig. 3e-f to check whether a similar AF correction worked for increasing IL concentrations. Fig. 4d shows the result of these experiments: the recovered ATTO 550 phase lifetime is properly recovered no matter what IL concentration is used (red curve), and readily distinguished from phantom AF (black curve). Since AF increases with IL concentration, the phase lifetime SDV increased (blue curve), as already observed for increasing phantom thicknesses (Fig. 3b).

In summary, the previous measurements show that in the presence of a fluorescence signal sufficiently larger than the phantom autofluorescence (independent from its thickness), reasonable estimation of the sample’s fluorescence lifetime can be performed. Moreover, a purely analytical uncorrelated background subtraction approach can be used, which can be useful when an AF measurement cannot be obtained, as might be the case for *in vivo* measurements.

### Live cell imaging behind phantoms

Having characterized the phase lifetime behavior of homogeneous fluorescent samples imaged through highly scattering medium, we used our time-gated approach to image a group of cells through similar phantom layers as a crude model for a tumor xenograft located subcutaneously. The basic question we asked was whether it is possible to distinguish unlabeled cells from cells expressing the fluorescent protein (FP) mCyRFP1^34^, with the goal of distinguishing cells targeted by a molecular marker (FP expressing cells) and untargeted cells.

Fig. 5 summarizes the result of these cell experiments. Figs. 5a-b, which shows the fluorescence images of FP-expressing cells (Fig. 5a) and control cells (Fig. 5b) in the absence of phantom, clearly illustrating the larger brightness of the FP-expressing cells compared to the control cells (estimated number of cells in the FOV: 300,000, excitation power: 12 mW, total integration time: 25 sec). Fig. 5c, by contrast, shows the fluorescence image of the FP-expressing cells imaged through a 1.5 mm thick phantom layer, with a major decrease in the fluorescence intensity (3 times lower). Unlike cells with no phantom layer above them, the fluorescence from these cells is spread all over the frame, as expected for a sample behind a phantom. Figs. 5e-f shows FP-expressing cells’ and the control cells’ phasors using the numerical ucB subtraction approach described earlier (both numerical ucB and phantom AF subtraction were tested in the mCyRFP1 experiments, resulting in very similar phase lifetimes, as shown in *Supporting information* Fig. S6B). Fig. 5g, which represent the phase lifetimes extracted from the different samples observed through different phantom thicknesses, clearly show that cells which expressed mCyRFP1 exhibit a constant lifetime for all thicknesses (∼3.3 ns), despite the decrease in their fluorescence intensity, as observed previously for the dye samples used in Figs. 2 & 4. By contrast, the control cells, characterized by a weak AF (2551 counts for cells with no phantom on top) barely distinguishable from that of the phantom, exhibited an increasing phase lifetime for larger phantom thickness, becoming indistinguishable from that of the phantom AF (∼3.5 ns), as expected for a weak fluorescence signal. Notice that the measured mCyRFP1 lifetime, while similar to the phantom’s AF lifetime, is clearly distinguishable for all phantom thicknesses, thanks to its larger fluorescence intensity. In the presence of a large amount of unlabeled cells however, as would be the case for a subcutaneous FP-expressing tumor surrounded by live tissue, this extrinsic fluorescent signal might become overwhelmed by cell AF and would therefore be detectable only in favorable conditions of quantum yield, labeled cell concentrations and tissue scattering properties, or by limiting the excited region to dimensions comparable to the labeled cells region.

## Discussion

Optical imaging of fluorescent dyes has emerged as a powerful imaging method in preclinical applications due to the fast, noninvasive nature and quantitative results achieved by this approach. Interest in fluorescence-based techniques for clinical applications remain high due to their potential to target and detect specific tissues with high sensitivity, providing a fast, noninvasive and quantitative readout. Due to the wide range of bright and biocompatible contrast agents and relatively low-cost instrumentation, the visible range of the electromagnetic spectrum would be attractive for clinical fluorescence-based imaging methods. However, a primary challenge with fluorescence based imaging in the visible range is the development of detection methods that will overcome the major attenuation of light intensity in this region due to the high scattering and absorption of the tissue. Here, we demonstrated that phasor-based fluorescence lifetime imaging using a highly sensitive time-gated SPAD camera is capable of detecting fluorescent dyes and proteins emitting in the visible spectrum through highly scattering tissue-like phantom. Despite the limitations of visible range imaging mentioned above, as well as the omnipresence of tissue autofluorescence, this work illustrates the potential of wide-field time-gated fluorescence lifetime imaging for this kind of application.

In particular, we showed that despite the presence of autofluorescence background, it is relatively straightforward to detect and identify the presence of a mass of FP-expressing cells characterized by a distinct lifetime behind up to 1.5 mm of phantom. FP have become a popular tool for live-cell and small animal *in vivo* imaging, in particular for the study of localization, motility and interaction of proteins in living cells ^35, 36^, mostly in the visible range. Their rapid rise as genetically manipulated imaging probes provides unique opportunities for real-time tracking of specific cells *in vivo*, enabling visualization of changes in target-gene promoter activity, tracking cellular movement in embryogenesis and inflammatory processes, monitoring migration of small parasites within a host, studying important aspects of cancer, such as tumor cell trafficking, invasion, metastasis and angiogenesis^37-40^. Yet, in the case of *in vivo* applications, optical imaging of intact tissue does not provide cellular or intra-cellular resolution and hence, like PET, is limited to macro or meso-localization of the fluorescent probe^41, 42^. Therefore, there is a continuing interest to develop new systems for whole-body imaging. Recent whole-animal imaging of FP-labeled tumors have been extended to NIR FP^43, 44^, the tumors in these studies being either subcutaneous or consisting of several million cells in deep tissue. The very low tissue AF observed in this region of this spectrum will allow the method reported here to achieve a similar goal. Thus, e.g., Shcheslavskiy *et. al*^*45*^ recently presented a macro-FLIM system for *in vivo* imaging purposes, with a relatively high spatial resolution and molecular specificity, enabling the detection of endogenous NAD(P)H fluorophores in tumor. They used a scanning TCSPC technique, which usually covers a relatively low scanning area, of 1×1 mm, but by placing the objects in the intermediate image plane of a confocal scanner, were able to achieve a larger FOV of more than 1 cm. Lyons *et. al* ^46^have developed a computational method based on a single-photon time-of-flight camera enabling imaging of an object embedded inside a strongly diffusive medium over more than 80 transport mean free paths in the NIR, paving a way toward a single photon based detectors, as SS2, to be used for real time, *in vivo* imaging. Extending the results described here to the NIR, in particular work involving the A549 cells transfected with mCyRFP1, which show FLI through highly scattering medium, will be our next step toward developing a new *in vivo* imaging method with improved contrast, sensitivity and specificity for *in vivo* detection of FP or fluorescently-labeled probes.

## Acknowledgments

This work was funded in part by NIH Grant GM 095904 and CRCC Grant CRR-18-523872 (UCLA), and in part by the Swiss National Science Foundation Grant 166289 and the Netherlands Organization for Scientific Research Project 13916. Rinat Ankri thanks Prof. Dror Fixler from the Faculty of Engineering at Bar Ilan University, Israel, for financial support to this work.

