## Supporting information for "Single Photon, Time-Gated, Phasor-based Fluorescence Lifetime Imaging Through Highly Scattering Medium"

**Content:**

1) Figures:

- Figure S1: Regions of interests (ROIs) that were used in phasor analysis.
- Figure S2: Time-gated decays of Cy3B sample behind various phantom thicknesses.
- Figure S3: Cy3B sample analysis.
- Figure S4: Phantom-only measurements.
- Figure S5: ATTO 550 measurements.
- Figure S6: Lifetime analysis of A549 expressing mCyRFP1 and control cells.

2) Data used in this paper.

**Figure S1: The regions of interests (ROIs) were used for the phasor transformation in Figures 2-5.** (a) Small 4x4 ROIs that were used for phasor transformation, while each ROI resulted with a single phasor point on the UC. (b) Large ROI that was used when calculating the mean fluorescence intensity of a given sample. Only the bottom part of the image was used for the calculation, since the upper part of the image presented higher dark counts values.

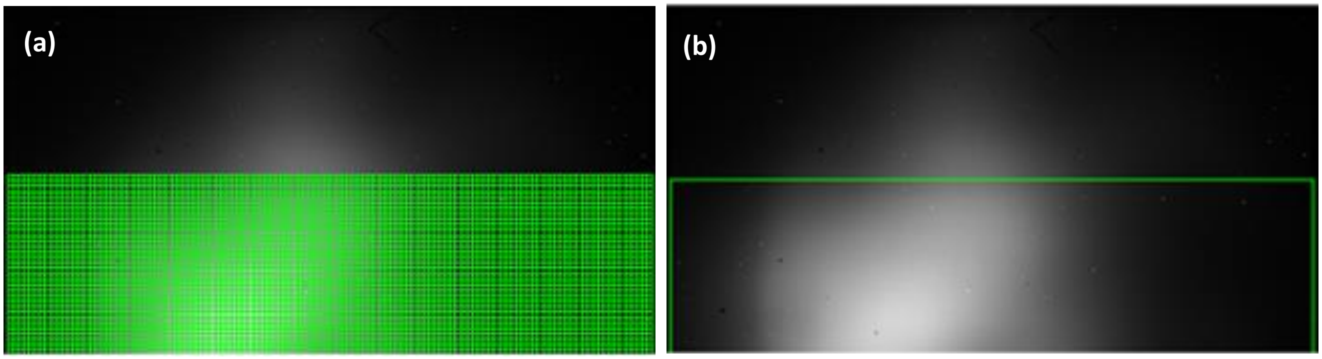

**Figure S2: Time-gated decays of Cy3B sample behind various phantom thicknesses.** a: Full-period (50 ns) representation of the gated signal (gate width: 15 ns, gate step: 428.6 ps, 117 gates) for the same Cy3B sample observed without phantom (black) or behind increasing phantom thicknesses. The decay corresponds to the sum of all pixels in the lower half of the datasets. b: Semi-logarithmic representation of the same decays, showing the quasi-overlap of the different decays.

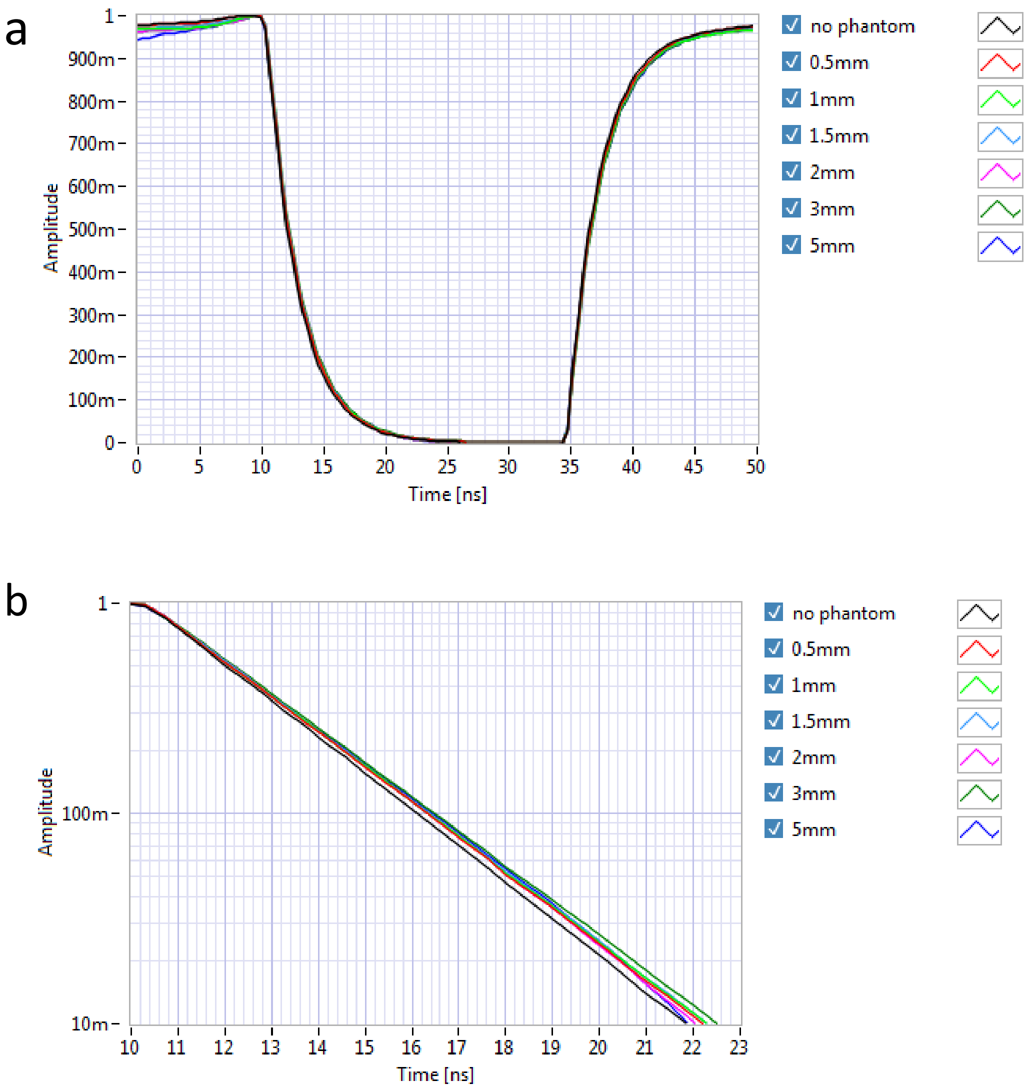

**Figure S3: Cy3B sample analysis.** a: Dependence of the fitted uncorrelated background on exposure time (black dots). A linear fit (red) is also shown. b: Dependence of the collected uncorrelated background-corrected fluorescence intensity on phantom thickness (black dots) and exponential fit (red). The calculated exponential length scale parameter is 0.77 ± 0.27 mm.

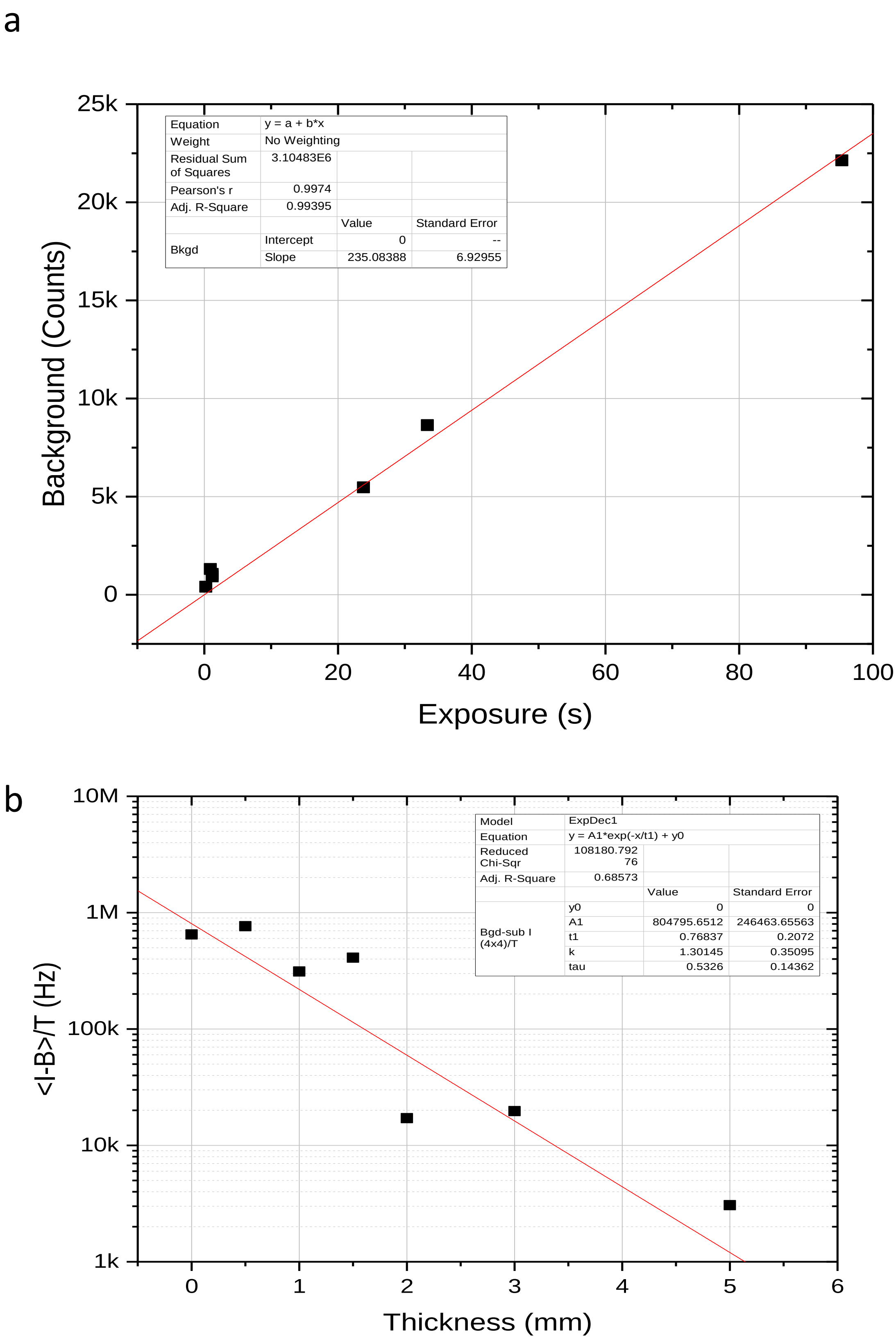

**Figure S4: Phantom-only measurements.** a: Background-subtracted count rate, indicating a fairly constant autofluorescence brightness of the phantoms. b: Fitted background rate from the same measurements. As expected for an uncorrelated background due to detector noise, this contribution is equal for all measurements.

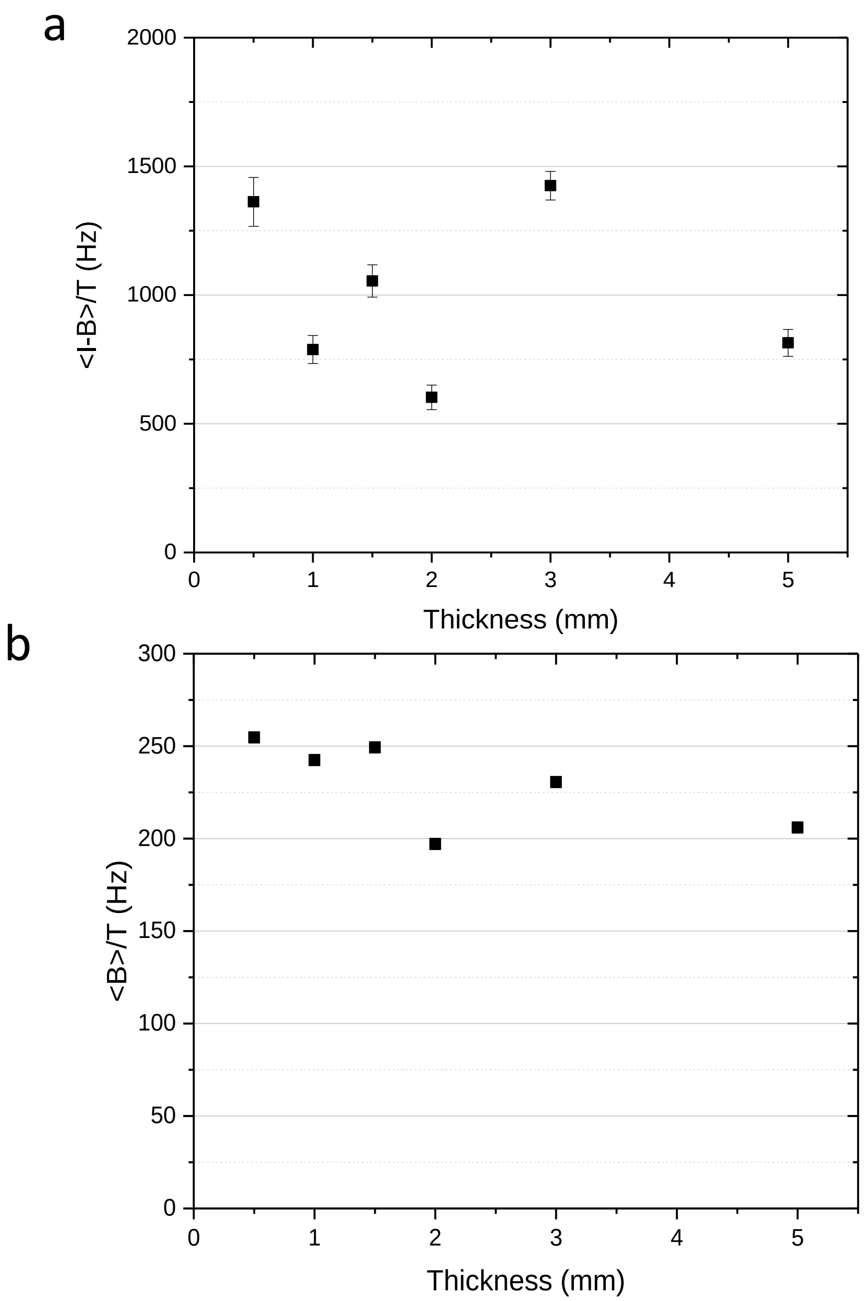

**Figure S5: ATTO 550 measurements.** a: Dependence of the collected uncorrelated background-corrected fluorescence intensity on phantom thickness (black dots) and exponential fit (red). The calculated exponential length scale parameter is 0.70 ± 13 mm. b: The measured phantom autofluorescence count rate (~ 1 kHz) can be used to estimate its contribution to the ATTO 550 signals studied in Fig. 4. Its contribution is steadily increasing with increasing thickness, reaching 58 % for the 5 mm phantom. The model fitted to the data is discussed in the text.

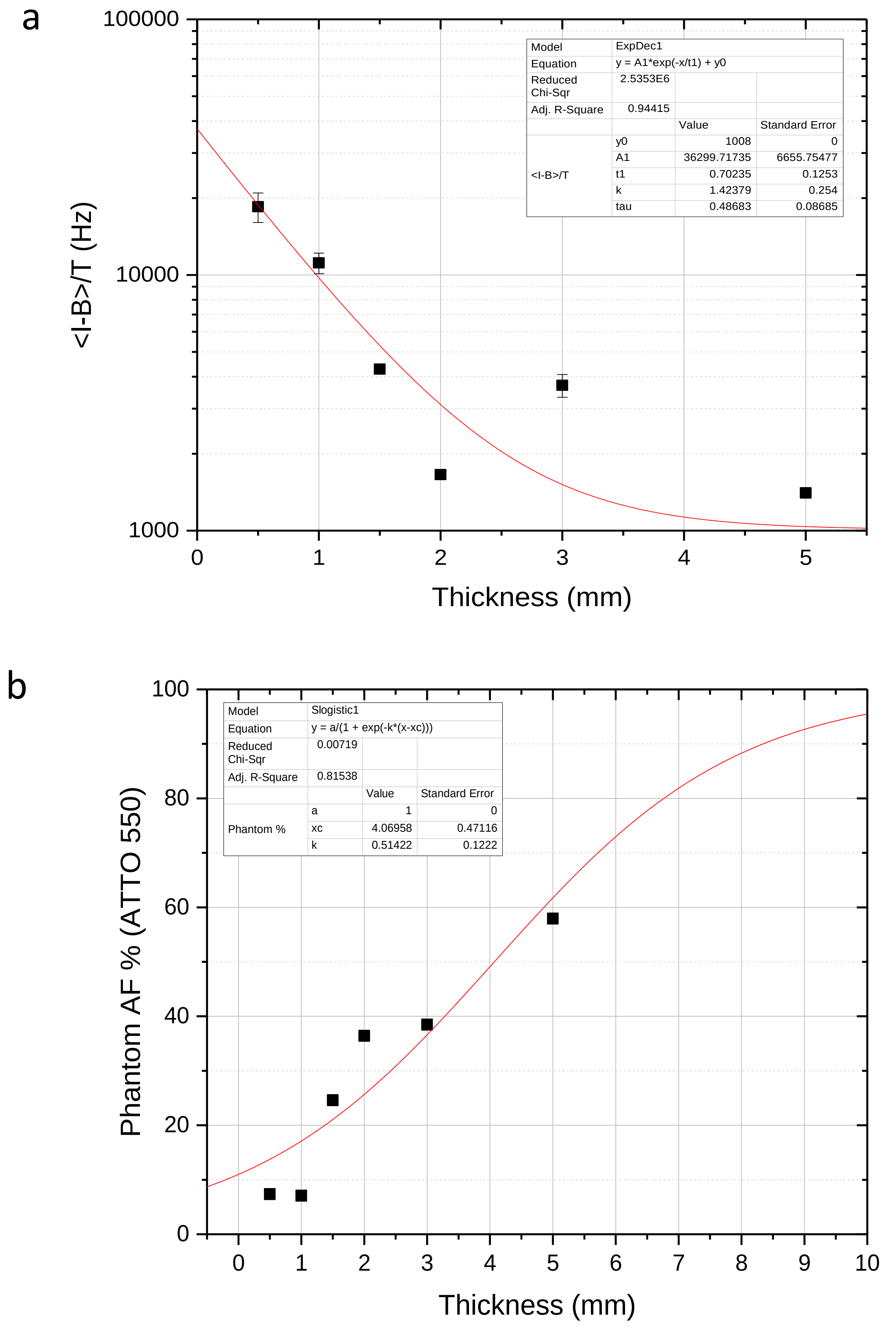

**Figure S6: Phase lifetime analysis of A549 cells expressing the fluorescent protein mCyRP1 behind phantom layers of 0.5, 1 and 1.5 mm in thickness, and for a sample with no phantom layer.** (a) The 4x4 pixels ROIs were used in the phasor transformation of the cells fluorescence. (b) The dashes line is the lifetime extracted following software background subtraction and the solid line is for lifetime extracted following phantom layer subtraction (same integration times for sample and background phantom, of 10 msec). Results suggest that there is not a significant change in the resulted lifetimes between the SW and the phantom BG correction (ATTO550 lifetime behind the 1.5 mm phantom, which looks different in values, presented τ=3.28 for the phantom BG correction and 3.31 for SW BG correction). (c) The fluorescence intensity values for mCyRFP1 transfected cells (dotted line), wild type cells (dashed line) and phantom layers (solid line). Intensity was measured for images following numerical BG subtraction. All data points are the mean intensity of the pixels within the ROI (presented in panel a)

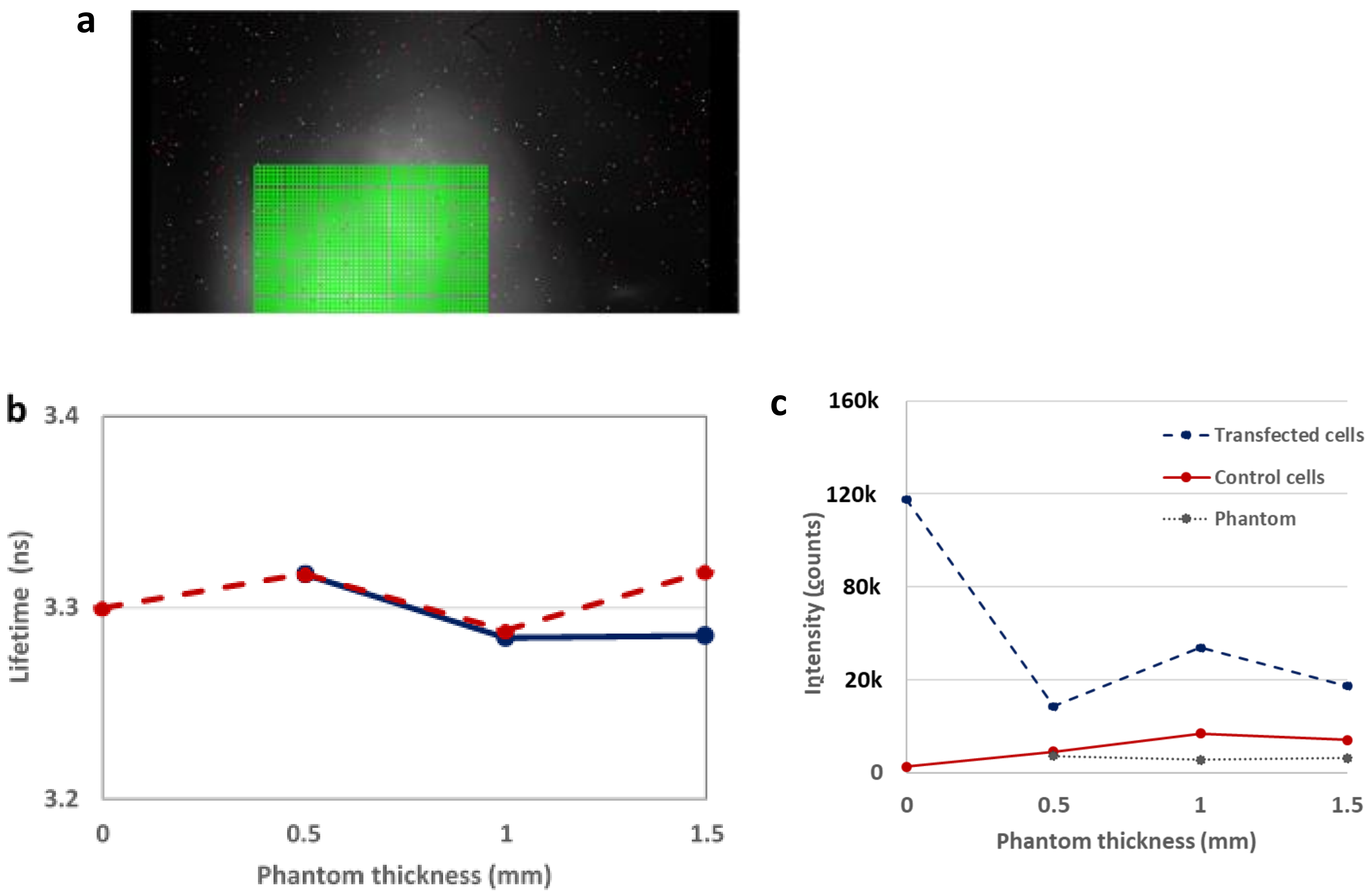

**Data used in this paper**

Supporting Data for; *Single Photon, Time-Gated, Phasor-based Fluorescence Lifetime Imaging Through Highly Scattering Medium* is available at *FigShare, at:* Ankri, R.; Basu, A.; Ulku, A. C.; Charbon, E.; Weiss, S.; Michalet, X., Supporting Data for Single Photon, Time-Gated, Phasor-based Fluorescence Lifetime Imaging Through Highly Scattering Medium: https://figshare.com/articles/Data_Single_Photon_Time-Gated_Phasor_based_Fluorescence_Lifetime_Imaging_Through_Highly_Scattering_Medium/8290199
